# Prediction and analysis of skin cancer progression using genomics profiles of patients

**DOI:** 10.1101/393454

**Authors:** Sherry Bhalla, Harpreet Kaur, Anjali Dhall, Gajendra P. S. Raghava

## Abstract

Metastatic state of the Skin Cutaneous Melanoma (SKCM) has led to high mortality rate worldwide. Previously, various studies have revealed the association of the metastatic melanoma with the diminished survival rate in comparison to primary tumors. Thus, prediction of melanoma at primary tumor state is crucial to employ optimal therapeutic strategy for prolonged survival of patients. The RNA, miRNA and methylation data of The Cancer Genome Atlas (TCGA) cohort of SKCM is comprehensively analysed to recognize key genomic features that can categorize various states of metastatic tumors from primary tumors with high precision. Subsequently, various prediction models were developed using filtered genomic features implementing various machine learning techniques to classify these primary tumors from metastatic tumors. The SVC model (with class weight and RBF kernel) developed using 17 mRNA features achieved maximum MCC 0.73 with sensitivity, specificity and accuracy 89.19%, 90.48% and 89.47% respectively on independent validation dataset. Our study reveals that gene expression based features performs better than features obtained from miRNA profiling and epigenomic profiling. Our analysis shows that the expression of genes *C7, MMP3, KRT14, KRT17, MASP1*, and miRNA hsa-mir-205 and hsa-mir-203a are among the key genomic features that may substantially contribute to the oncogenesis of melanoma even on the basis of simple expression threshold. The major prediction models and analysis modules to predict metastatic and primary tumor samples of SKCM are available from a webserver, CancerSPP (http://webs.iiitd.edu.in/raghava/cancerspp/).

## Introduction

Cancer is one of the major causes of mortality worldwide since last few decades. According to GLOBOCAN, 2012, 14.1 million new cancer cases and 8.2 million deaths has been estimated worldwide (Lindsey A. Torre MSPH Freddie Bray PhD Rebecca L. Siegel MPH Jacques Ferlay ME Joannie Lortet☐Tieulent MSc Ahmedin Jemal DVM, 2015). Among them, skin cancer is one of the most commonly recognized cancers in the United States. Evidently, an estimation of 91,270 melanoma new cases and 9,320 deaths have been reported in 2018. Additionally, it is more prominent in females than males (Society, 2018). The malignant transformation of normal human epithelial melanocytes which are located within the basement membrane of the skin results in melanoma development. There are several genetic and environmental factors contributes to melanoma carcinogenesis, which include excessive exposure of UV radiations, indoor tanning devices and contacts with certain chemical like arsenic and hydrocarbons etc (Schaeken and van der Hoeven, 1990).

Recently, with the advancement of genomic technologies, there is a huge increment in the generation of big genomic data, particularly in the field of cancer (Ana-Teresa Maia, 2017) which can be explored for the identification of diagnostic and prognostic cancer biomarkers (Meyerson et al., 2010). The cancer genome atlas (TCGA) project is one of the leading comprehensive repository associated with the genomic, epigenomic, transcriptional profiles/gene expression, DNA methylation, clinical data and mutational data includes somatic variations, copy number variation (CNV), somatic copy number alterations (SCNA) (Tomczak et al., 2015). The TCGA study has revealed the four subtypes of SKCM based on mutant genes such as mutant BRAF, mutant RAS, mutant NF1, triple WT (wild-type) (Cancer Genome Atlas, 2015). Further, it has been observed that the mutational rate of these genes is much higher in melanoma patients than other cancer samples of TCGA. Interestingly, over 50% of melanoma patients have BRAF kinase (B-Raf proto-oncogene, serine/threonine kinase) mutations (Guan et al., 2015). In addition, various studies have demonstrated that skin cutaneous melanoma arises from the anomalies in genetic and epigenetic factors such as expression of mRNAs and miRNAs, aberration in methylation patterns of CpG islands of genes and histone modifications, which paves the way for development of potential molecular biomarkers in melanoma (Hoon et al., 2004;Mori et al., 2005;Mori et al., 2006;Tanemura et al., 2009;Zheng et al., 2009;Goto et al., 2010;Philippidou et al., 2010;Schinke et al., 2010;Kanemaru et al., 2011;Mazar et al., 2011a;Mazar et al., 2011b;Nguyen et al., 2011;Greenberg et al., 2014). In past, several reports reveal the potential role of microRNA expression as prognostic biomarkers in cutaneous melanoma. For instance, miR205 and miR29c both act as tumor suppressors and down-regulate the expression of E2F1, E2F5 (Dar et al., 2011) and DNMT3 (Nguyen et al., 2011) genes respectively‥ Besides miRNAs, Histone methyltransferases also act as crucial players in the progression of melanoma by enhancing the expression of zeste homolog 2 (EZH2) (Tiffen et al., 2015).

Earlier studies have scrutinized the distinctions between primary melanoma and metastatic melanoma (Hoek et al., 2004;Haqq et al., 2005;Smith et al., 2005;Talantov et al., 2005;Mischiati et al., 2006;Winnepenninckx et al., 2006;Jensen et al., 2007). The metastasis mechanism involved several pathways including epithelial mesenchymal transition (EMT), angiogenesis and invasion (Geiger and Peeper, 2009). Furthermore, the aggressive stage of melanoma can metastasize to lymph nodes, distinct tissues and organs (Geiger and Peeper, 2009). Although the survival of cutaneous melanoma patients affected by various factors, the disease’s early stage diagnosis is one of the most vital parameter with the greatest impact on survival. Different studies have shown the metastases free MM patients *i.e.* patients with primary tumor have significantly prolonged survival (White et al., 2002;Soong et al., 2010). Evidently, the five year overall survival rate of melanoma patients *i.e.* 20%, 60% and 90% for distinct, regional stage and localized tumors respectively (Society, 2018). Hence, the detection of tumor at localized stage *i.e.* primary tumor is crucial for patient management and implementation of appropriate therapeutic strategy for prolonged survival of patients. Previously, several stochastic stage wise prediction and classification methods have been developed for diverse cancer types (Jagga and Gupta, 2014;Bhalla et al., 2017). Recently, one of the studies has distinguished primary and metastatic SKCM samples and identified important miRNA and RNA expression based putative biomarkers for progression of SKCM metastasis (Li et al., 2015). They also found the association of metastatic progression score with various clinical characteristics like Breslow’s depth, Clark’s level etc. The lack of gold standard performance measures, performance on independent validation dataset and webserver to analyse new data based on those identified markers are the major lacuna. The current study is an attempt to overcome that inadequacy. In this analysis, we have made an effort to understand the cutaneous skin melanoma progression based on TCGA data include RNA-seq, miRNA-seq and methylation expression; we develop prediction model on the basis of various machine learning techniques that can segregate primary and metastasized skin cutaneous melanoma (SKCM) patients. In this study, we have also made an attempt to develop prediction model for categorization of different types of metastatic states from primary tumor.

## Material and methods

### Datasets

RNA-seq, miRNA-seq and methylation profiling data for human skin cutaneous melanoma (SKCM) retrieved from The Cancer Genome Atlas (TCGA) project using TCGA - Assembler 2 (Wei et al., 2017). In addition, manifest, biospecimen files and files containing clinical information such as new tumor events, drugs, age, gender etc. were also downloaded to extract clinical parameters using Biospecimen Core Resource (BCR) IDs of patients/subjects. Finally, we obtained 466 patients [102 primary tumor, 74 Regional Cutaneous or Subcutaneous Tissue (includes satellite and in-transit metastasis), 222 Regional Lymph Node and 68 Distant Metastasis samples] for RNA and methylation data, whereas 444 samples available [95 primary tumor, 64 Regional Cutaneous or Subcutaneous tissue (includes satellite and in-transit metastasis), 214 Regional Lymph Node and 71 Distant Metastasis] for miRNA expression data. We referred these primary tumor, Regional Cutaneous or Subcutaneous Tissue (includes satellite and in-transit metastasis), Regional Lymph Node and Distant Metastasis samples as P1, P2, M1 and M2 respectively. Clinical characteristics of these patients displayed in **Figure S1**.

In present study we have used RNA and miRNA expression profiles in terms of RSEM values for 20,502 genes and 1,870 miRNAs respectively. We downloaded the methylation profiles for 20,879 genes (acquired using the Illumina Human-Methylation450K DNA Analysis BeadChip assay, based on genotyping of bisulfite-converted genomic DNA at individual CpG-sites). Notable, we have downloaded all CpG sites for each gene no matter whether they are DNAse hypersensitive or not. This provides Beta values, a quantitative measure of DNA methylation (Bibikova and Fan, 2009; Bibikova, et al., 2006; Du, et al., 2010). Here, methylation value for each gene represents the mean of methylation beta values of all CpG sites located on each individual gene.

### Pre-processing of Data

#### Normalization of miRNA and RNA Expression

It has been observed that there is a wide range of variation in RSEM values of RNAs and miRNAs, thus we transformed these values using log2 after addition of 1.0 as a constant number to each of RSEM value. Further, features with low variance have been excluded from the data using *caret* package in R, followed by z-score normalization of data. Thus, log2-transformed RSEM values for each mRNA and miRNA were centred and scaled by employing *caret* package in R. Following equations were used for computing the transformation and normalization:

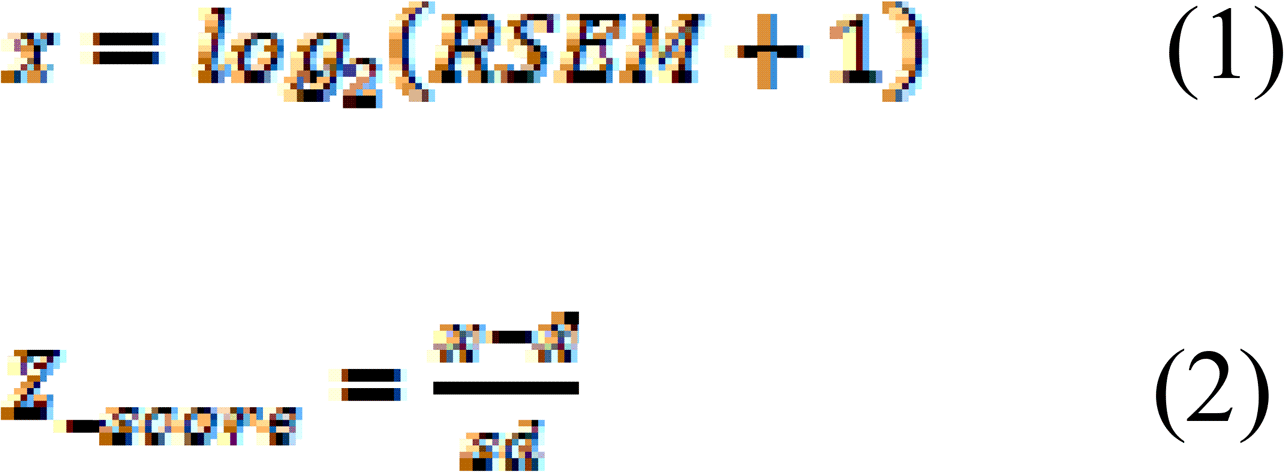

Where *Z__scoτe_* is the normalized score, *x* is the log-transformed, *x̄* expression and is the mean of expression and *sd* is the standard deviation of expression.

It has been also observed that in some of patients, RNA expression available for both tissue and blood samples. Here we have taken the average of both the samples for each patient.

#### Feature Selection Techniques

One of challenges in developing prediction model is to extract important features from the large dimension of features. In this study, we implemented two techniques *i.e.* SVC with L1 penalty employing Scikit package (Fabian Pedregosa, 2010) and ‘SymmetricalUncertAttributeSetEval’ with search method of ‘FCBFSearch’ of WEKA software package (Smith and Frank, 2016). We filtered genes (RNA expression), methylation pattern of genes and miRNA expression as features that can distinguish metastatic samples from primary tumor samples using these techniques. The FCBF (Fast Correlation-Based Feature) algorithm employed correlation to identify relevant features in high-dimensional datasets in small feature space (Lei Yu 2003). SVC-L1 method selects the non-zero coefficients and then applies L1 penalty to select relevant features to reduce dimensions of the data. To select the robust features, first the data was split into the ratio of 80:20 for 10 times followed by features selection using SVC-L1 or WEKA on each occasion from the training dataset. From this resampling process, we obtained 10 sub-sets of features. We have selected the subset having highest performance. To check the robustness of features, we computed the average stability index (Jaccard index) using OmicsMarkeR package (Determan, 2015) for each subset and finally reported the overall stability index. For all the signatures the average stability index is nearly in the range of 0.40 to 0.43.

#### Implementation of machine learning techniques

Firstly, we have developed the prediction models to categorize primary tumor and metastatic samples based on selected genomic features using various classifiers implementing Scikit package. These classifiers include ExtraTrees (Wehenkel, 2006), KNN, Random forest (Ho, 1998), Logistic Regression (LR) (Lin, 2011), Ridge classifier and SVC - RBF (radial basis function) kernel with class weight factor were implemented employing scikit package (Fabian Pedregosa, 2010). In addition, to understand the progression of skin cancer from primary tumor to metastasis, we also analyse and develop prediction models using various machine learning classifiers of scikit package based on genomic features (RNA expression) to classify the sub-categories of metastatic samples from primary tumor samples *i.e.* Intralymphatic tumors ((P2) v/s primary tumor (P1), lymphatic tumors (M1) v/s primary tumors (P1), distant metastatic tumors (M2) v/s primary tumors (P1), regional (P1P2) v/s lymphatic tumors (M1) and metastatic tumors (M1M2) v/s regional tumor (P1P2).

The optimization of the parameters for the various classifiers was done by using grid search with PR curve as scoring performance measure for selecting best parameter. As our data is imbalanced, it is very important to select the parameters that show balanced precision and recall.

#### Visualization of samples

After applying supervise learning to classify samples, we visualised the distribution of samples based on selected features on reduced dimensions using t-SNE (t-Distributed Stochastic Neighbour Embedding) methods implementing the two R Packages; Rtsne and scatterplot3D packages. t-SNE is a non-linear dimensionality reduction algorithm employed to analyze the high-dimensional data. It converts multi-dimensional data to two or more dimensions (Maaten, 2014).

#### Identification of important features using simple threshold based approach

Here, we employed MCC (Matthews correlation coefficient) based feature selection technique to identify important features and developed single feature based prediction models to distinguish metastatic samples from primary tumor samples. Single feature based models are also called threshold based models in which feature having a score below a specific threshold is assigned to metastatic tumor if it is downregulated in metastatic tumor samples otherwise as primary tumor sample and vice versa. We compute performance of each given feature and identify features having highest performance in term of MCC with minimum difference in sensitivity and specificity.

#### Performance Evaluation of Models

In present study, both internal and external validation techniques were employed to evaluate the performance of models. Here, primarily main dataset has been subdivided into two subsets *i.e.* training dataset and independent or external validation dataset in ratio of 80:20. We used 80% of the main dataset for training and remaining 20% for independent validation. First, the training dataset is used for developing model and for performing ten-fold cross validation as internal validation. In this ten-fold cross validation technique, training dataset is randomly splits into ten sets; of which nine out of ten sets were used as training sets and the remaining tenth set as testing dataset. This process is repeated ten times in such a way that each set is exploited once for testing. The final performance of model is the mean performance of all the ten sets. In order to avoid optimization of parameters in case of tenfold cross-validation, we also implement external validation. In case of external validation, we evaluate our model on an independent or external dataset not used for training. In order to measure performance of models, we used standard parameters. Both threshold-dependent and threshold-independent parameters were employed to measure the performance. In case of threshold-dependent parameters, we measure sensitivity, specificity, accuracy and Matthew’s correlation coefficient (MCC) using following equations.

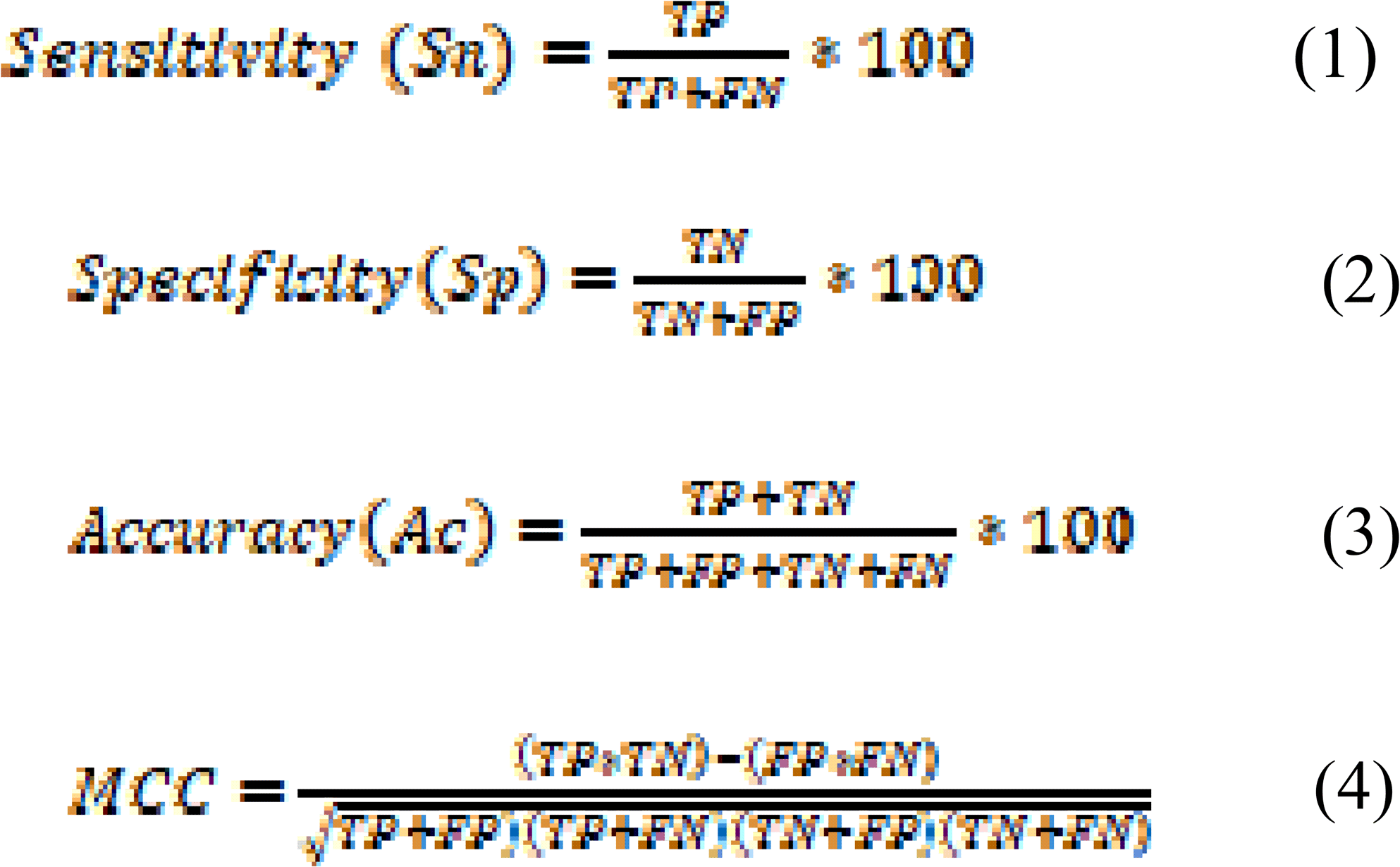

Where, FP, FN, TP and TN are false positive, false negative true positive and true negative predictions respectively.

While, for threshold-independent measures, we used standard parameter Area under the Receiver Operating Characteristic curve (AUROC). The AUROC curve is generated by plotting sensitivity or true positive rate against the false positive rate (1-specificity) at various thresholds. Finally, the area under AUROC curve calculated to compute a single parameter called AUROC.

#### Functional annotation of signature genomic markers

In order to discern the biological relevance of the signature genes, enrichment analysis is performed using Enrichr (Kuleshov et al., 2016). Enrichr executes Fisher exact test to identify enrichment score. It provides Z-score and adjusted p-value which is derived by applying correction on a Fisher Exact test. Further to understand the biological impact of miRNAs in metastatic melanoma development, we employed miRTarBase (Chou et al., 2018) to identify target genes of signature miRNA.

## Results

In current study, we have analysed the RNA-seq, miRNA-seq, methylation-seq data of SKCM from TCGA for 466, 444, 466 patients respectively, to mine important features which can discriminate various degree of metastatic tumors (P2, M1 and M2) from the primary tumors (P1) using machine learning techniques. The pipeline depicting the course of this study is shown in Figure 1.

**Figure1:**
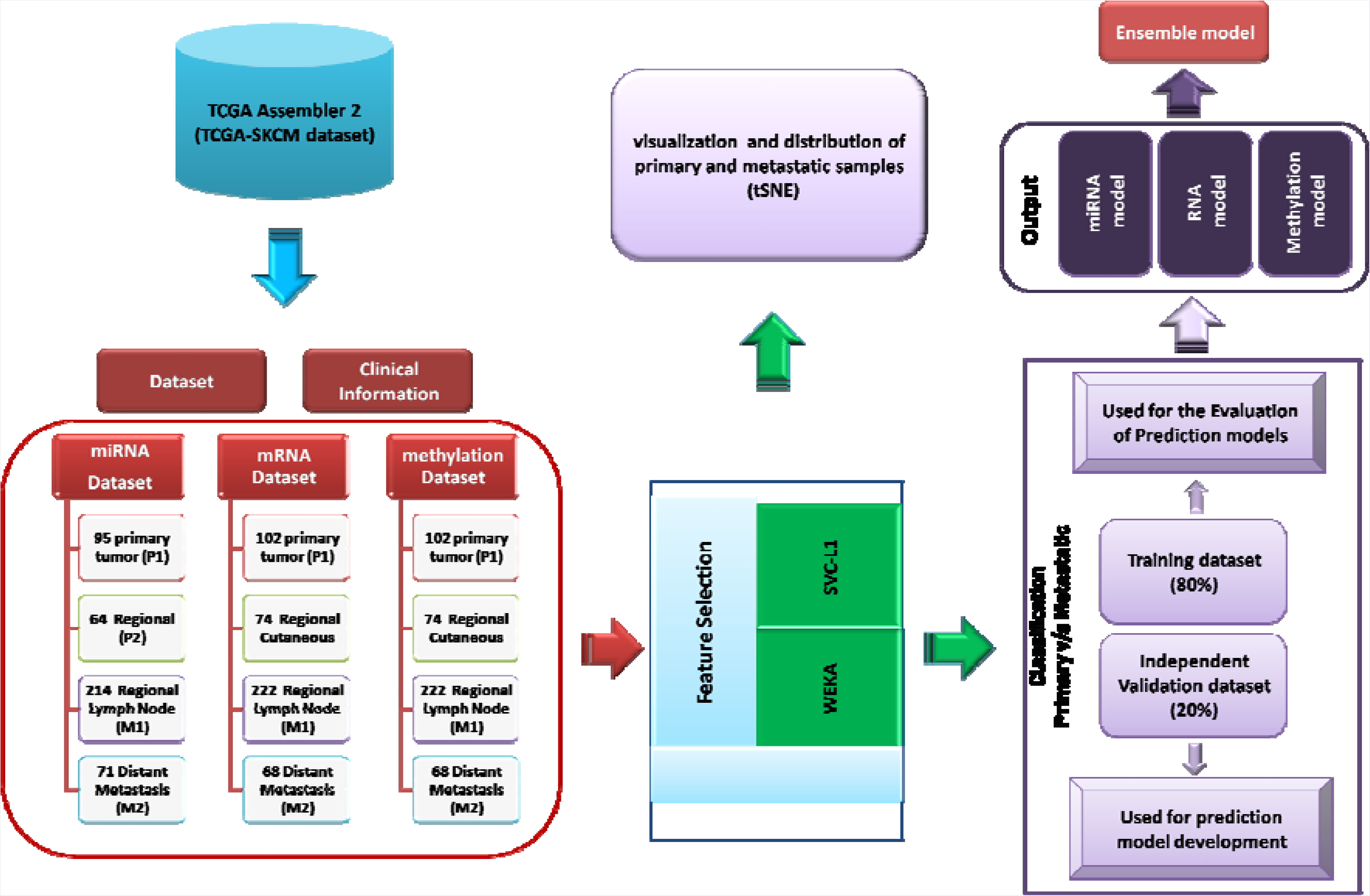
Pipeline to represent the workflow of the study.

## Gene Expression based models

With aim to classify metastatic and primary tumor samples, firstly RNAseq data of 466 patients containing expression of 20,502 genes was used to select relevant features using two feature selection methods SVC-L1 and WEKA-FCBF. We obtained nearly 140 and 17 features using WEKA-FCBF and SVC-L1 respectively. We applied six machine learning algorithms on the selected features obtained using above two methods. As shown in Table 1, nearly 92.76% (sensitivity) metastatic tumors and 90.12% (specificity) primary tumors of training dataset and 89.19% metastatic and 90.48% primary tumor samples of validation dataset are correctly identified by SVC-model based on these 17 features (selected by SVC-L1). This model achieved accuracy 92.18% and 89.47% with MCC of 0.79 and 0.73 on training and validation dataset respectively. The boxplot depicting the expression pattern of these 17 features in metastatic and primary tumor samples is shown in Figure 2. Interestingly, model based on 140 features selected using WEKA-FCBF attained similar performance (data not shown), but as the number of features are quite high as compared to features selected in SVC-L1, we preferred and reported the model on 17 features.

**Table 1:**
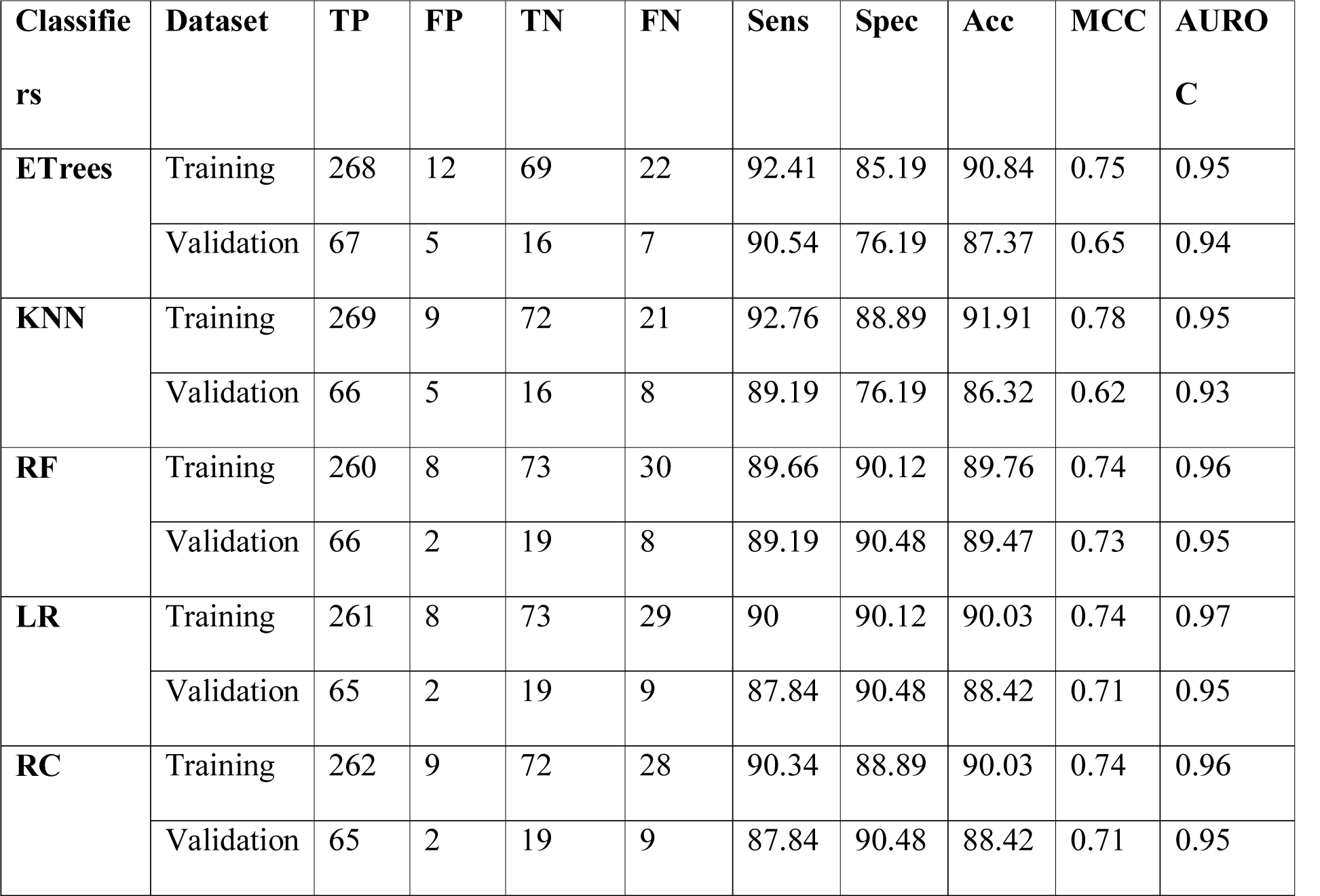
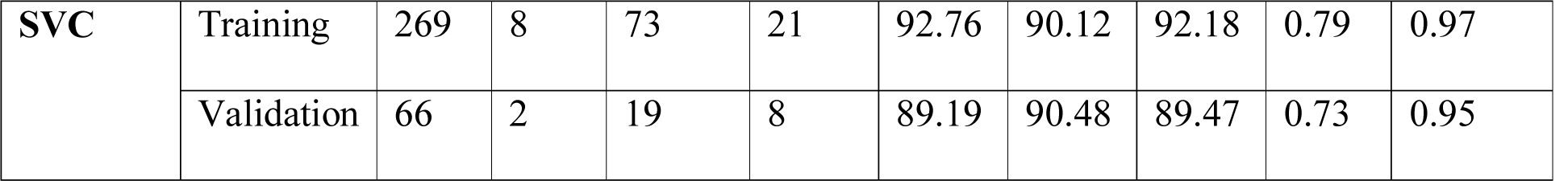
Performance measures of 17 RNA expression based features (selected by SVC-L1 feature selection method) on training and independent validation dataset to classify metastatic from primary tumor samples applying various machine-learning algorithms (classifiers).

| **Classifiers** | **Dataset** | **TP** | **FP** | **TN** | **FN** | **Sens** | **Spec** | **Acc** | **MCC** | **AUROC** |
| --- | --- | --- | --- | --- | --- | --- | --- | --- | --- | --- |
| **ETrees** | Training | 268 | 12 | 69 | 22 | 92.41 | 85.19 | 90.84 | 0.75 | 0.95 |
|  | Validation | 67 | 5 | 16 | 7 | 90.54 | 76.19 | 87.37 | 0.65 | 0.94 |
| **KNN** | Training | 269 | 9 | 72 | 21 | 92.76 | 88.89 | 91.91 | 0.78 | 0.95 |
|  | Validation | 66 | 5 | 16 | 8 | 89.19 | 76.19 | 86.32 | 0.62 | 0.93 |
| **RF** | Training | 260 | 8 | 73 | 30 | 89.66 | 90.12 | 89.76 | 0.74 | 0.96 |
|  | Validation | 66 | 2 | 19 | 8 | 89.19 | 90.48 | 89.47 | 0.73 | 0.95 |
| **LR** | Training | 261 | 8 | 73 | 29 | 90 | 90.12 | 90.03 | 0.74 | 0.97 |
|  | Validation | 65 | 2 | 19 | 9 | 87.84 | 90.48 | 88.42 | 0.71 | 0.95 |
| **RC** | Training | 262 | 9 | 72 | 28 | 90.34 | 88.89 | 90.03 | 0.74 | 0.96 |
|  | Validation | 65 | 2 | 19 | 9 | 87.84 | 90.48 | 88.42 | 0.71 | 0.95 |
| **SVC** | Training | 269 | 8 | 73 | 21 | 92.76 | 90.12 | 92.18 | 0.79 | 0.97 |
|  | Validation | 66 | 2 | 19 | 8 | 89.19 | 90.48 | 89.47 | 0.73 | 0.95 |

**Figure 2:**
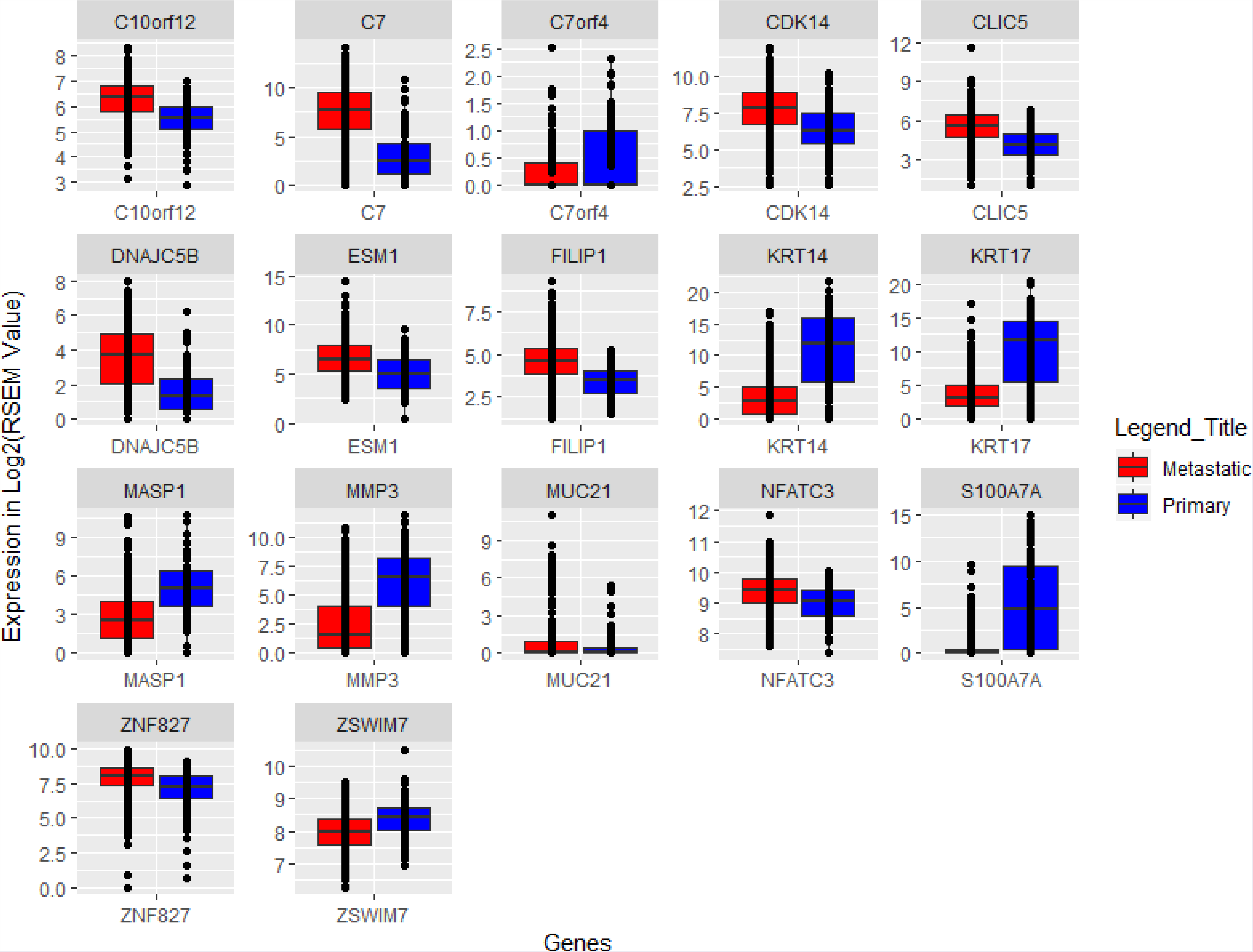
Boxplot representing the expression pattern of 17 genes selected using SVC-L1.

As we analysed the new tumour event (NTE) clinical file of SKCM patients, we observed that 16 patients with primary tumor have been shown to be in distant metastasis with new tumour events. Therefore, we remove or exclude these 16 samples from dataset and again developed the classification model. There was marginal increase of MCC from 0.73 to 0.77 on validation dataset (Supplementary Table S1).

Enrichment analysis shows the biological role of the RNA signature in carcinogenesis. Evidently out of 17 signature features (genes), *C7* and *MASP1* are involved in Complement system activation (p-value < 0.05), while *KRT17* and *KRT14* are part of intermediate filament component (p-value < 0.05). It has been shown that metastatic cancer cells use actin bundles to disrupt from a primary tumour and invade the surrounding tissue. After travelling in the vasculature or lymphatic system, they exit into a new niche and form a new tumour (Hanahan and Weinberg, 2011).

## miRNA expression based models

Next, miRNA expression explored to elucidate its role in progression of metastasis in SKCM. The number of miRNA features selected by WEKA-FCBF and SVC-L1 is 32 and 5 features respectively. SVC-W model based on 32 miRNAs attains maximum performance with MCC of 0.69 on training dataset and 0.66 on validation dataset. Further, nearly 86.33% metastatic samples and 90.79% primary samples of training dataset and 90.14% metastatic samples and 78.95% primary samples of validation dataset were correctly predicted (**Table 2**). The mean expression pattern of these 32 miRNA in primary and metastatic samples is represented in **Supplementary Figure S2**. The Logistic Regression model based on 5 miRNAs (feature selected by SVC-L1 method), achieved maximum MCC of 0.62 on training dataset and 0.59 on validation dataset. This model correctly predicted 86.69% metastatic samples and 81.58% primary tumor samples of training dataset and 83.1% metastatic samples and 84.21% primary samples of validation dataset (**Table 3**). These 5 miRNAs include hsa-mir-205, hsa-mir-218.2, hsa-mir-513a.1, hsa-mir-675 and hsa-mir-7974.

**Table 2:**
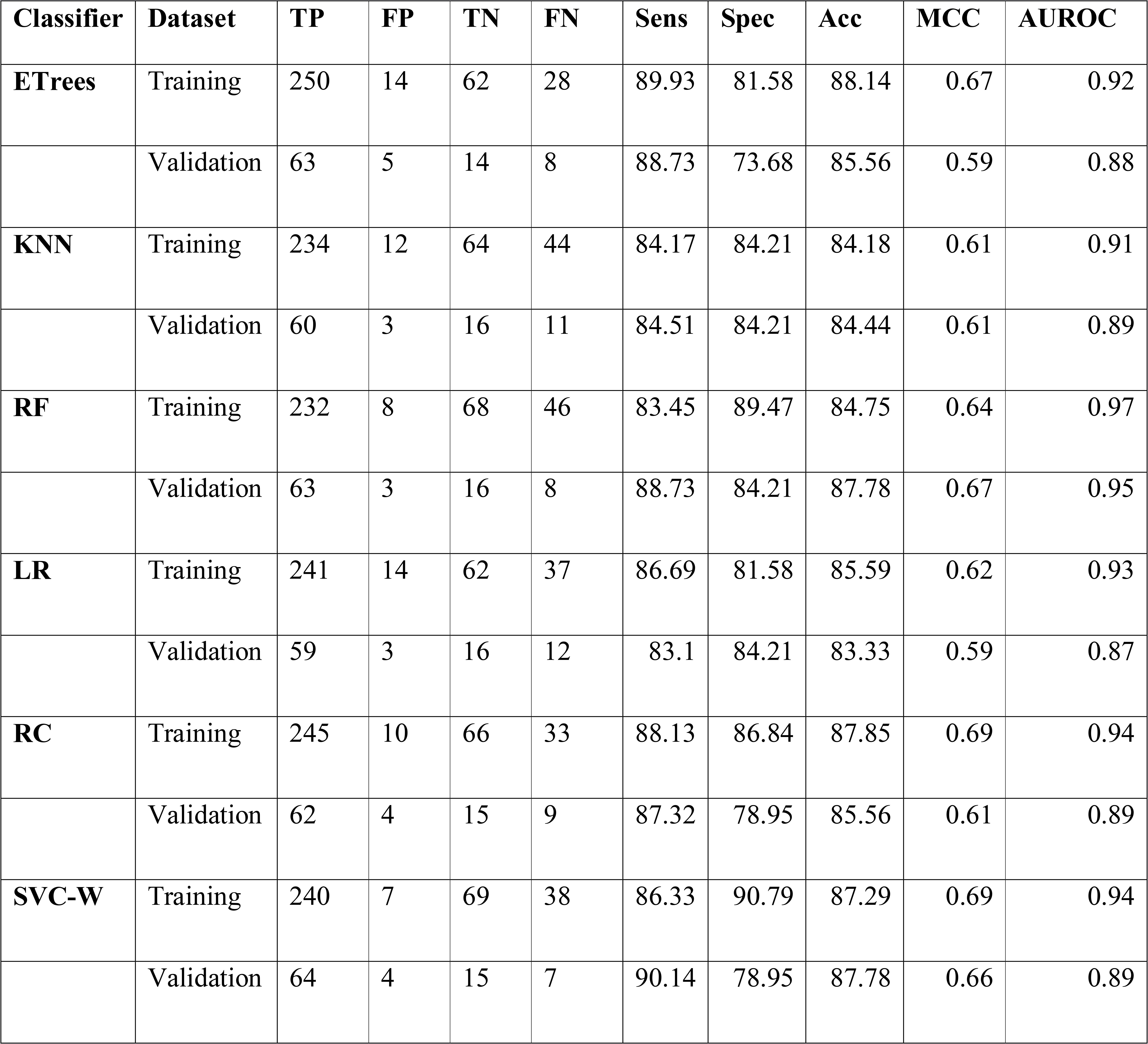
Performance measures of 32 miRNA expression features (selected by WEKA-FCBF feature selection method) on training and independent validation to classify metastatic from primary samples dataset by applying various machine-learning algorithms.

**Table 3:**
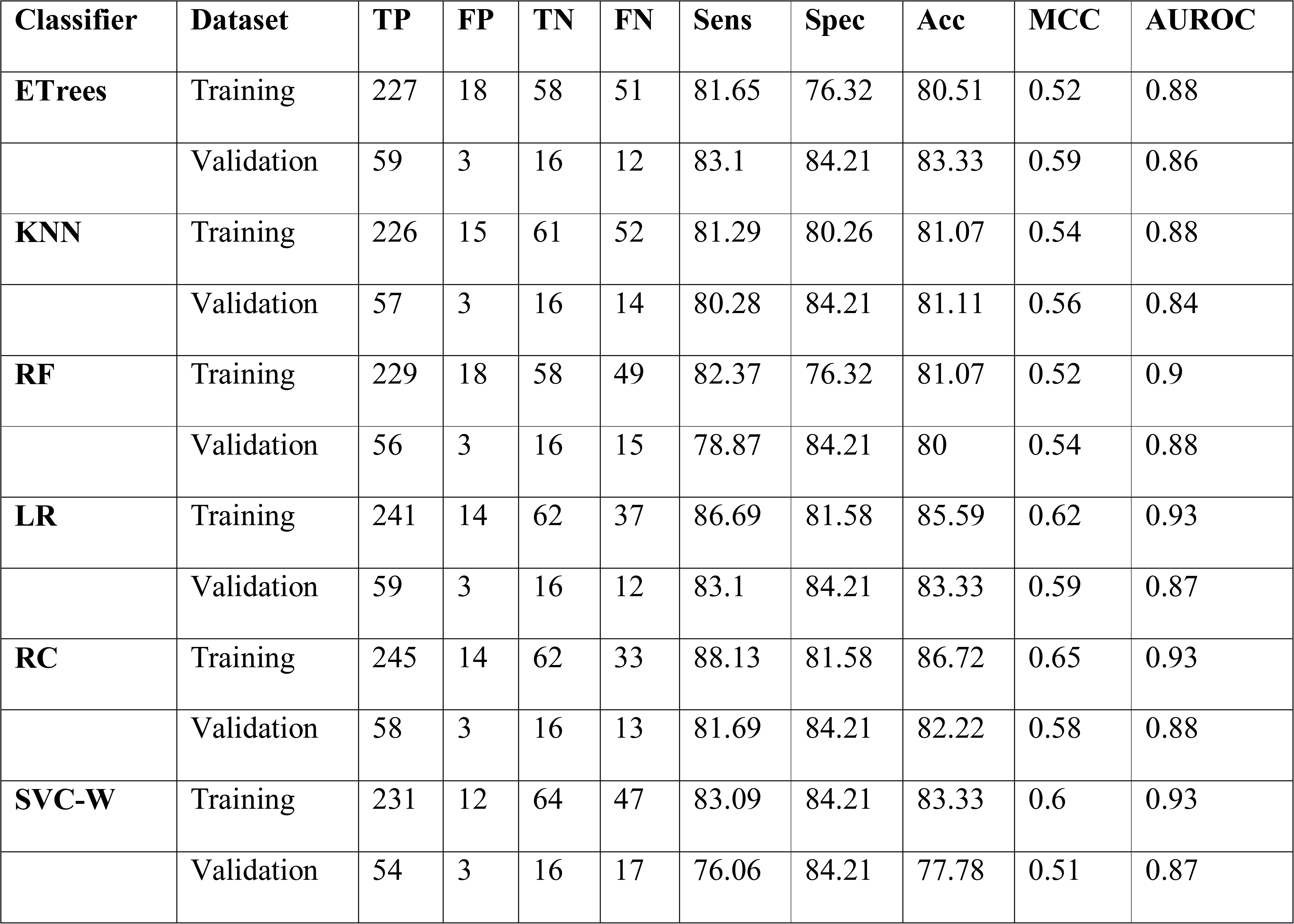
Performance measures of 5 miRNA expression features (selected by SVC-L1 feature selection method) on training and independent validation dataset to classify metastatic from primary samples by applying various machine-learning algorithms.

Among these signatures, hsa-mir-205 targets various genes (identified from miRTarBase) such as *ZEB2, ZEB1, ERBB3, PRKC, ERBB2, E2F1, BCL2, ITGA5, VEGFA, AR, SMAD4, EGLN2, ERBB2, BCL2, E2F1, LAMC1, VEGFA, SMAD1, SRC, VEGFA, DDX5* and *YES1* etc. Gene enrichment analysis have shown that these genes are significantly enriched in various cell growth promoting and oncogenesis associated pathways including transcriptional misregulation, TGF-beta signaling, wnt signaling, PDGF signaling, EGFR signaling, PI3K signaling, p53 signaling, ErbB signaling, VEGF signaling, cell cycle, hypoxia and angiogenesis, apoptosis processes etc. This analysis signifies the role of hsa-mir-205 as tumor suppressor in melanoma development as it gets downregulated with the progression of metastatic melanoma.

## Methylation based model

To ascertain the role of epigenetics in progression of melanoma from primary to metastatic, we have taken average methylation value for each gene as described in methods. Firstly, 38 and 2 features were selected using WEKA-FCBF and SVC-L1 respectively. Subsequently, classification models were developed using 38 features and it can be observed (in Table 4) that average methylation values are not very good predictors to distinguish metastatic and primary tumor samples. For instance, LR model based on these 38 features achieved maximum performance, able to discriminate them with maximum MCC of 0.48 and 0.44 on training and validation dataset respectively. It correctly predicted only 76.47% metastatic samples and 79.27% primary tumor samples of training dataset and 78.38% of metastatic samples and 71.43 % primary tumor samples of validation dataset (Table 4).

**Table 4:**
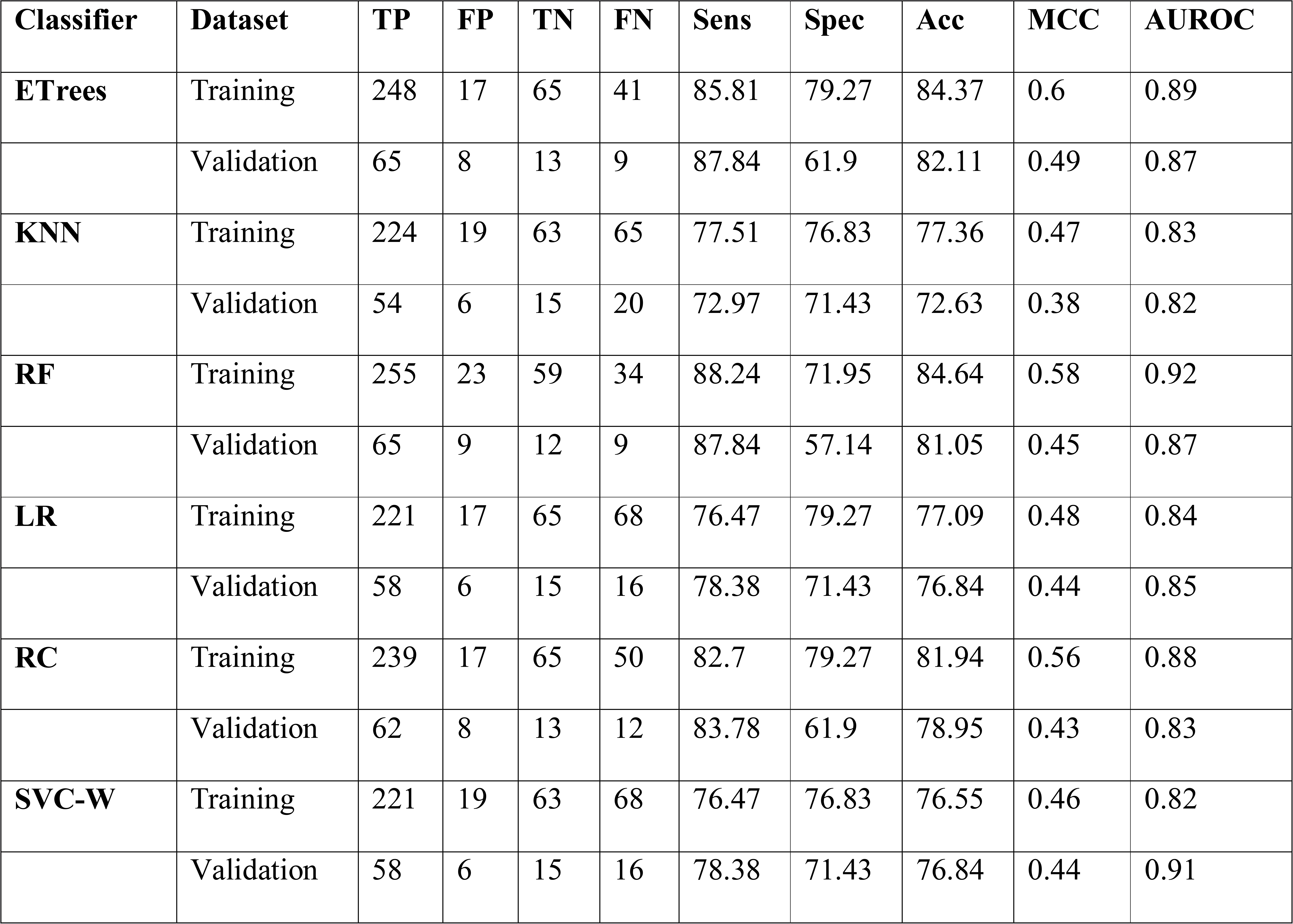
Performance measures of 38 features or average methylation of genes (features selected using WEKA-FCBF feature selection method) on training and independent validation dataset to classify metastatic from primary samples by applying various machine-learning algorithms.

### Visualization of samples

As we observed that RNA expression based features are performing quite well as compared to miRNA expression and methylation profiling, we visualised the 17 RNA expression features using tSNE. The Figure 3 shows that primary tumors (P1) form the separate cluster (red colour) in comparison to different states of metastasis. Substantial number of P2 samples differ from P1 but some of them merges/co-clustered with P1 which is quite expected as P1 progresses to P2 (Figure 3A). The tSNE analysis shows a clear distinction between P1 and M1 (Figure 3B) with some of the primary samples going extreme into the boundaries of M1. Surprisingly the distant metastatic samples are quite widely distributed in comparison to Primary tumors as shown in Figure 3C. Finally the visualization of these seventeen features in three dimensional space in all the four classes is shown in (Figure 3E). These analyses prompt us to develop specific prediction models for classifying each state of metastasis from primary samples.

**Figure 3:**
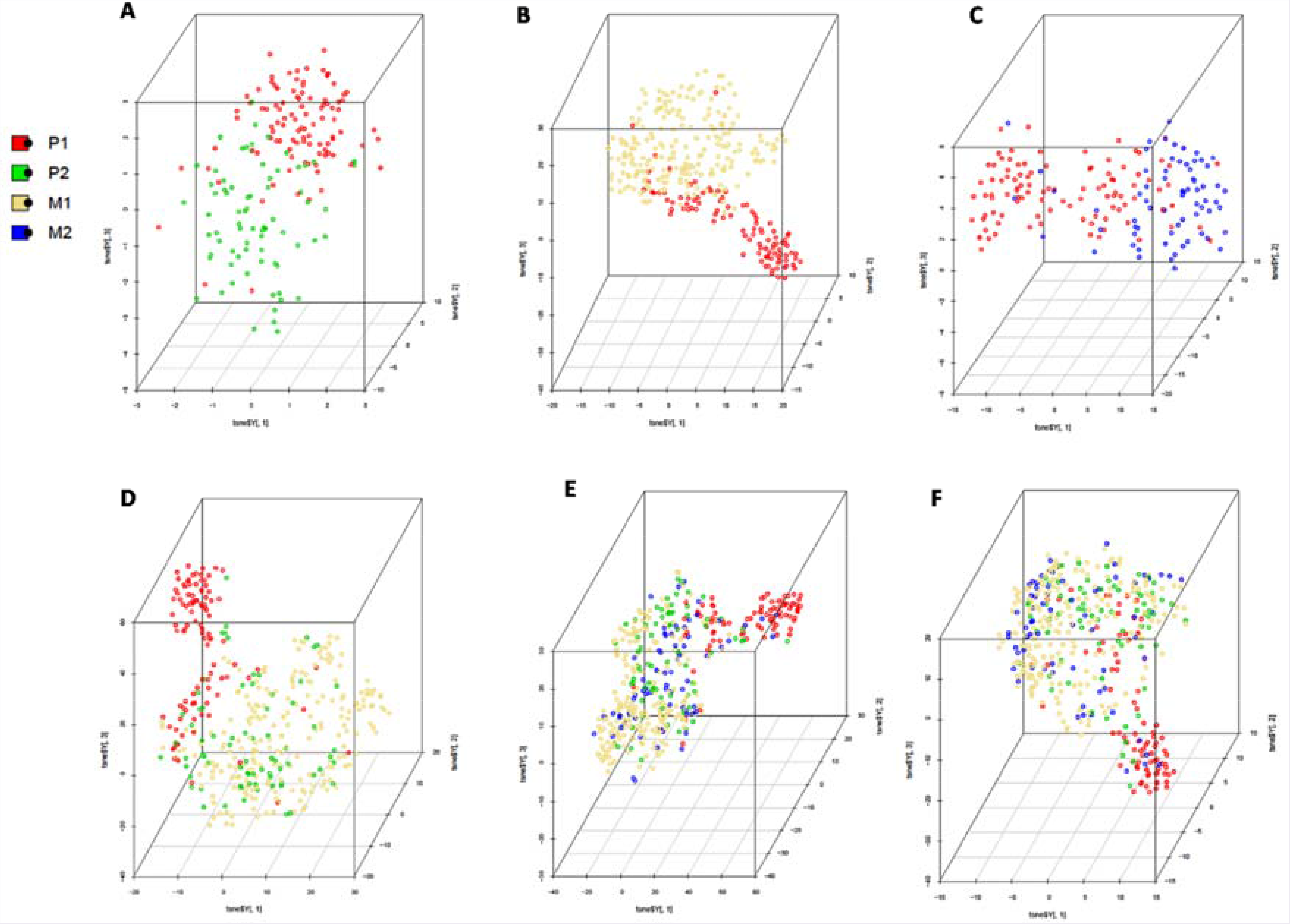
Scatterplot3D view of tSNE dimension reduction of 17 selected features: (A) distribution of P1 and P2 samples on 17 features; (B) distribution of P1 and M1 samples;(C) distribution of P1 and M2 samples; (D) distribution of primary tumors (P1) in comparison to in transit and satellite tumors (P2) from lymphatic metastatic tumors (M1); (E) distribution of P1, P2, P3 and P4 samples; (F) distribution of P1, P2, P3 and P4 samples after removing 16 primary tumor samples (observed as distant metastatic in NTE file).

## Ensemble Model

Next, in order to compile information from individual models developed using all the three types of genomic features, we developed ensemble method. In the ensemble method, prediction score from each model *i.e.* RNA, miRNA and methylation were provided as input features to SVC. This model attained MCC of 0.73 and 0.71 on training and validation dataset respectively (Table 5).

**Table 5:** Performance measures of RNA-seq, miRNA-seq and methylation-seq ensemble features on training and independent validation dataset to classify metastatic from primary samples by applying SVC.

| **Dataset** | **TP** | **FP** | **TN** | **FN** | **Sens** | **Spec** | **Acc** | **MCC** | **AUROC** |
| --- | --- | --- | --- | --- | --- | --- | --- | --- | --- |
| **Training** | 244 | 5 | 71 | 34 | 87.77 | 93.42 | 88.98 | 0.73 | 0.97 |
| **Validation** | 60 | 1 | 18 | 10 | 85.71 | 94.74 | 87.64 | 0.71 | 0.93 |

## Discrimination between Primary and sub-categories (or various states) of metastasis

### Intra-lymphatic tumors v/s primary tumors

The primary tumors (P1) are localised lesions and P2 includes the samples with in transit metastasis and satellite metastasis which represents intra-lymphatic tumour which has not still spread to lymphatic nodes. We selected 10 features using SVC-L1 (as in the above models RNA-seq data selected appropriate number of features using SVC-L1) and the results of classification models on these 10 features shows that it is difficult to classify the samples with intra lymphatic tumour (P2) from primary tumour (P1). The KNN-based model correctly identified metastatic and primary patients of training data is 84.75% and 93.83% with MCC of 0.79 and the maximum correctly identified metastatic and primary patients is 73.33% and 85.71% with MCC 0.60 of validation dataset (Supplementary Table S2). The selected ten features is shown in column 1 of heatmap (Figure 4).

**Figure 4:**
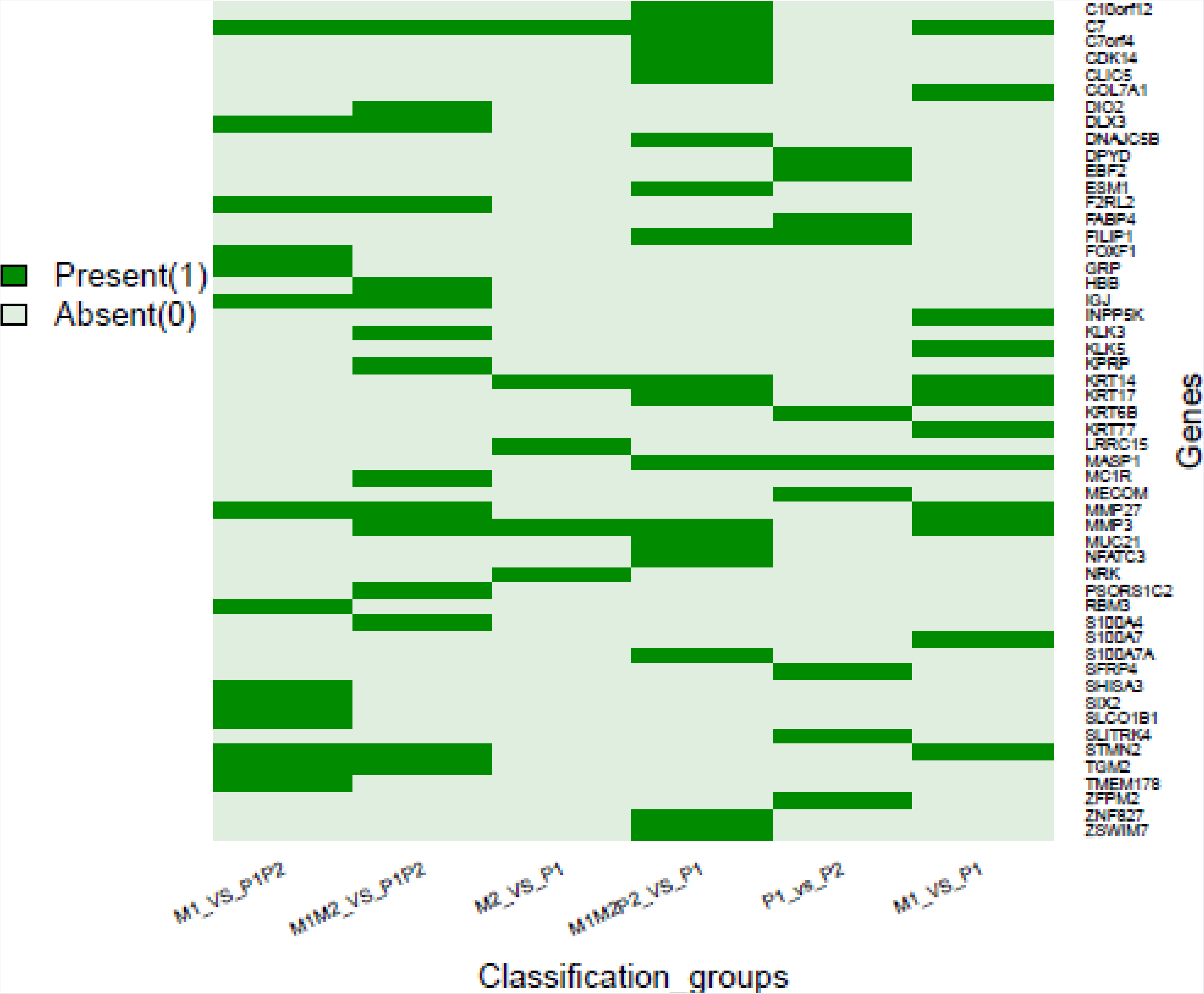
Heatmap showing presence and absence of various features in different signature sets developed for segregating metastatic samples from primary samples.

### Lymphatic tumors v/s primary tumors

Further we tried to classify tumors that have invaded lymphatic nodes from primary tumors. Our analysis shows these tumors can be classified with high precision. As SVC-W based model based on 12 genes (Figure 4) selected using SVC-L1 feature selection method distinguished samples with good sensitivity of 97.74%, specificity 91.36% and MCC of 0.90 of training data. We also observed the fair sensitivity of 95.56% and specificity of 90.48% along with MCC of 0.86 on validation dataset. This indicates that once the tumour has reached the lymph nodes, there is substantial variation in the expression of genes associated with metastasis in comparison to the primary or localized tumor (Table 6).

**Table 6:**
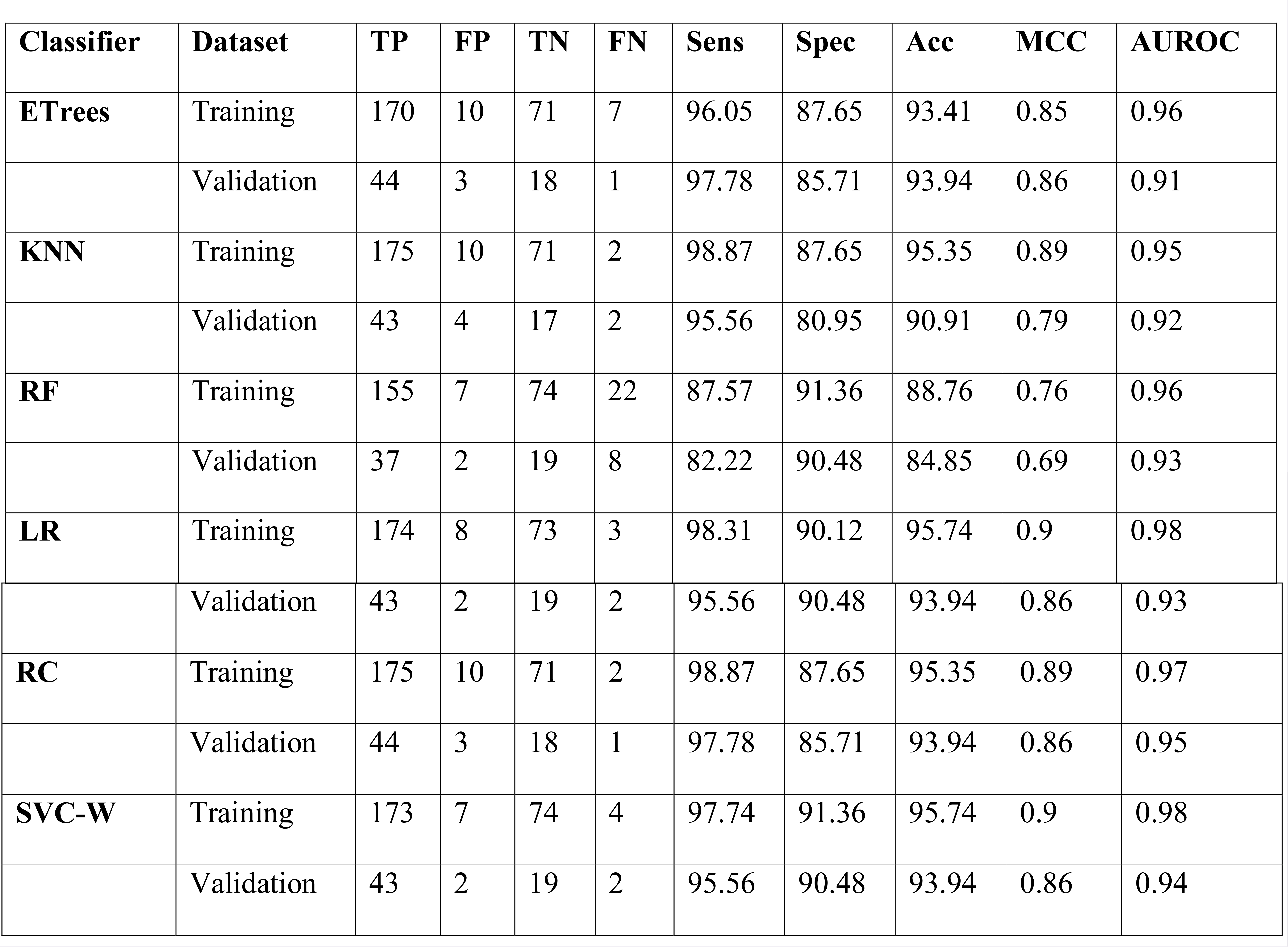
Performance measures of 12 RNA expression features (selected using SVC-L1 feature selection method) to discriminate M1 from P1 on training and independent validation dataset by applying various machine-learning algorithms.

### Distant Metastatic tumors v/s primary tumors

Next, we tried to classify the distant metastatic tumors (M2) from primary tumors (P1). Surprisingly classification of these two groups of samples is not as good as lymphatic node v/s primary on 5 features (Figure 4). The KNN model correctly classified 87.04% distant metastatic samples and 92.59% primary samples with MCC of 0.80 of training data, 78.57% distant metastatic samples and 85.71% primary samples with MCC of 0.64 of validation dataset (Supplementary Table S3).

### Regional v/s Lymphatic tumors

To differentiate between the tumors which have spread to lymph nodes (M1) and regional tumors, we combined primary (P1) and in transit and satellite tumors (P2). LR model based on 14 features selected by SVC-L1, achieved the sensitivity of 92.09% and specificity of 90% with MCC of 0.82 on training data and the sensitivity of 93.33% and specificity of 83.33% with MCC of 0.78 on validation dataset (Supplementary Table S4).

### Metastatic tumors v/s regional or primary tumor

Further, we developed model to categorize the tumors which spread to lymph nodes or metastasized (M1 and M2) from the tumors which were localized (P1). The Logistic Regression (LR) model based on 15 features (Figure 4) correctly classified 89.36% metastatic samples of training dataset with MCC of 0.74 and 80.56% metastatic sample of validation dataset with MCC of 0.61 (Supplementary Table S5).

The Figure 4 shows the various signatures that are associated for discrimination primary tumors from various states of metastatic tumors. Out of these 52 genes 5 genes (*KRT17;COL7A1;KRT14;KLK5;S100A7*) are enriched in skin epidermis formation (p-value=0.001). *GRP* and *KLK5* are involved in positive regulation of phospholipase C-activating G-protein coupled receptor signaling pathway (p-value 0.03). Three genes *MMP3, KLK5* and *KLK3* are involved in regulating glutamic-type endopeptidase activity (p-value 0.02).

### Single feature based classifìcation model using RNA and miRNA expression

Our goal is to develop single feature based classification models to segregate metastatic and primary tumor samples. Towards this, we used each miRNA and mRNA to develop a simple threshold-based model to segregate metastatic and primary tumor samples. In threshold based model, a sample is classified as metastatic if the log2 RSEM value of feature (in feature is upregulated in metastatic) is higher than a threshold value otherwise it is primary sample. In these models, threshold is varied incrementally from minimum to maximum RESM value. Finally, that threshold is selected which have maximum MCC in classifying metastatic and primary tumor samples. Consequently all the RNA and miRNA sites are ranked on the basis of maximum MCC and minimum difference in sensitivity and specificity to assess the ability of each feature to classify metastatic and primary samples (Supplementary Table S6). Table S6 represents 20 mRNAs and 2 miRNA that can distinguish two types of samples with high precision. hsa-mir-205 and hsa-mir-203b are top 2 miRNAs that can classify metastatic and primary tumor samples with MCC with MCC 0.59 and 0.54 at thresholds log2 (RESM values) 4 and 1 respectively. Both of these miRNA are downregulated in metastatic samples which indicates their potential role in tumor suppression. *C7, ANXA8L2, ODZ2, KRTDAP* and *KRT14* are among top 5 mRNA that can discriminate metastatic and primary tumor samples with MCC 0.55, 0.52, 0.5, 0.62 and 0.59 at thresholds log2 RESM values 4, 2, 4, 4 and 9 respectively. Among them *C7* is upregulated in metastatic samples that reveals its role as oncogenesis in melanoma, while rest of genes observed as downregulated in metastatic that indicates their role in tumor suppression.

Furthermore this analysis indicates if the log2 (RSEM) value of *C7* in a sample is greater than threshold 4 then sample belongs to metastatic otherwise primary sample as it is upregulated in metastatic samples. If the log2 (RSEM) value of *KRTDAP* in a sample is smaller than threshold 4 then sample belongs to metastatic otherwise primary sample as it is downregulated in metastatic samples

### Combo or hybrid model

As from the above analysis we can observe that 17 RNAs and miRNA hsa-mir-205 have performed best for discriminating primary and metastatic tumours. Therefore we combined them and developed models using various machine learning algorithms (Supplementary Table S7). Performance of this hybrid model is almost similar with marginal increase in specificity in comparison to the model based on 17 RNA features.

### Web Server Implementation

To contribute the scientific community, we developed a web server, CancerSPP (Skin Cancer Progression Prediction). CancerSPP is designed for the analysis and prediction of metastatic and primary tumor of SKCM from RNAseq, miRNA and methylation expression data. The web server has two modules; Prediction module and Data analysis module.

### Prediction module

This module permit the users to predict different states of metastatic samples from primary tumor samples *i.e.* Intra-lymphatic tumors ((P2) v/s Primary tumor (P1), Lymphatic tumors (M1) v/s Primary tumors (P1), Distant Metastatic tumors (M2) v/s Primary tumors (P1), Regional (P1P2) v/s Lymphatic tumors (M1) and Metastatic tumors (M1M2) v/s Regional tumor (P1P2) utilizing RSEM expression quantification values of signature genes. The user needs to submit RSEM value of signature genes for every melanoma patient. In the input file, the number of patients represents the number of columns in the file. The output file is based on prediction strategy. Furthermore, the user can choose desired model for the prediction of melanoma samples.

### Data Analysis Module

This module is used to evaluate the role of each gene in various melanoma stages/states such as primary, regional metastatic, lymph node metastatic and distinct metastatic based on RNA expression profiles. Further, it provides the p-value for each feature and analyses the appropriate difference between primary and metastatic stages for RNA expression data. Moreover, it also incorporates threshold-based MCC of each feature and means expression values for the RNA-seq expression data in the distinct primary and metastatic state of SKCM.

### Discussion

There is emergence of synergising clinical and molecular profiles of tumors for predictive modelling in cancer biology to study tumor detection and progression. This prediction helps the physicians in making a suitable decision about the treatment course (McCarthy et al., 2004;Cruz and Wishart, 2007). Previous studies on SKCM have focussed on determining its sub types (Cancer Genome Atlas, 2015) and survival (Haqq et al., 2005;Winnepenninckx et al., 2006). Li *et al* have tried to discriminate the primary and metastatic tumors by repeating their algorithm independently multiple times and proposed the proportion of runs in which an SKCM tumor was assigned to the metastatic group as an index of that tumor’s metastatic progression. Although, they provide a score for metastatic expression; but the gold standard performance measures and the classification models are not available to the public (Li et al., 2015).

In present study we have identified important features and their classification potential using different machine learning algorithms for segregating metastatic tumors from primary tumors. Our analysis shows that RNA expression profiling is the strongest predictor of metastasis than miRNA expression profiling and methylation profiling. Some of RNA expression signatures identified in present study have been implicated in skin cancer. Furthermore, *C7* shown to be a potential tumor silencer gene and its expression is highly downregulated in various carcinomas such as ovarian cancer and non-small cell lung cancer (NSCLC) (Ying et al., 2016). In current study, this gene alone can correctly predict 86.54% metastatic samples with MCC of 0.55. Another gene *i.e. MMP3*, which has repeatedly observed in our different signatures of metastasis have been reported to acts as melanoma suppressor gene (McCawley et al., 2008). Further, previously *KRT14* a keratin gene, has shown to be downregulated in case of skin cancer (Alam et al., 2011). In our analysis, it classified metastatic and primary samples with high sensitivity 94.23% and low specificity 60% with overall MCC of 0.59. As the number of RNA signatures for classifying different states of metastasis (Figure 4) overlap with each other, reveals the important genes in progression of SKCM.

Beside signature RNAs, the expression of hsa-mir-205 among the miRNAs have shown to be downregulated in various solid tumors (Hulf et al., 2011). In our analysis, hsa-mir-205 alone can discriminates the metastatic and primary tumors with the sensitivity and specificity of 85.67% and 78.95% respectively with MCC of 0.59 if its expression is less than log2 (RSEM value) of 4. In recent past, it has been observed that hsa-miR-205 targets oncogenes such as *E2F1* and *E2F5* and downregulates their expression results in inhibition of melanoma cell proliferation (Dar et al., 2011). In addition, it also acts as tumor suppressor microRNA in skin carcinoma (Philippidou et al., 2010;Xu et al., 2012) breast cancer (Iorio et al., 2009) and prostate cancer (Hulf et al., 2013).

SVC-model based on the expression of 17 RNAs is best performer in discriminating metastatic samples from primary tumor samples. This model attained the sensitivity 89.19% (correctly predicted metastatic samples) and specificity 90.48% (correctly predicted primary tumor samples) with MCC of 0.73 on validation datasets. Furthermore, individual models based on RNA expression also developed which can differentiate primary tumors from each state of metastasis. Interestingly, it has been observed that primary tumor can be easily distinguished from tumors which have metastasized to lymph nodes; evidently prediction model which separates these two groups is top performer among all the combinations. Eventually, we assume that this study would be helpful to recognize these important genomic signatures in classification of primary tumor from metastatic tumor of SKCM. This may delineate towards their role in progression from primary to metastatic state of SKCM. Finally, we have developed the webserver CancerSPP to integrate all the prediction models and tools established in current study. CancerSPP can analyze the gene expression data of a sample and predict whether it is a primary tumor or metastatic with a score using RSEM values derived from RNA-seq and miRNA-seq and methylation beta values.

## Contributions

S.B. and H.K. collected the data and created the datasets, developed classification programs, implemented algorithms. S.B., H.K. and A.D. created the back-end server and front-end user interface. S.B., H.K., A.D. and G.P.S.R. analyzed the results. S.B., H.K. and A.D. wrote the manuscript. G.P.S.R. conceived and coordinated the project, helped in the interpretation and analysis of data, refined the drafted manuscript and gave complete supervision to the project. All of the authors read and approved the final manuscript.

## Acknowledgement

The authors acknowledge funding agencies J. C. Bose National Fellowship (DST). S.B., H.K., and A.D. are thankful to ICMR, CSIR and DST INSPIRE for providing fellowships.

**Abbreviations**
Etrees: Extra Trees Classifier
KNN: K Neighbors Classifier
RF: Random Forest
LR: Logistic Regression
RC: Ridge Classifier
SVC: Support Vector Machine
SVC-W: Support Vector Machine with weight factor
TP: True positive
FP: False Positive
TN: True Negative
FN: False Negative
Sens: Sensitivity
Spec: Specificity
Acc: Accuracy
MCC: Mathews Correlation Coefficient
AUROC: Area under Receiver operator curve References

